# Reference-based variant detection with varseek

**DOI:** 10.1101/2025.09.03.674039

**Authors:** Joseph M Rich, Laura Luebbert, Delaney K Sullivan, Reginaldo Rosa, Lior Pachter

## Abstract

Variant detection from sequencing data is fundamental for genomics and is the first step in a wide range of applications, ranging from genome-wide association studies to disease diagnosis. Widely used tools for variant detection utilize a *de novo* approach that is based on a combination of read mapping algorithms and statistical methods for identifying genetic variation from error-prone sequencing data. This approach has been successful, although the detection of insertion and deletion variants, as well as the detection of variants from low-coverage data, remain challenging problems. We introduce varseek, a reference-based approach to variant detection that provides large improvements in performance in these challenging cases. The varseek approach utilizes a k-mer pseudoalignment approach, which provides the ability to identify variants at single-cell resolution in single-cell transcriptomics data. We showcase the versatility and performance of varseek for detecting tumor-specific COSMIC variants in glioblastoma single-cell sequencing.

## 1 Introduction

The identification of genetic variants from sequencing reads with respect to a reference genome (variant calling) is a key step in many bioinformatics workflows. Traditional variant calling involves aligning sequencing reads to a reference genome, followed by quality control and statistical testing to determine positions in the reference genome where a collection of reads can be ascertained to differ from the reference genome (1–6). Genetic variants offer critical insights into both evolutionary biology and disease mechanisms, with many common conditions, such as Alzheimer’s disease, cardiovascular disease, and cancer—harboring variants associated with pathogenicity or susceptibility (7–11).

Cancer diagnostics and treatment, in particular, is an important application of variant calling, as there may be heterogeneous factors driving the disease that are unique to each individual and depend on specific mutations (12). The ability to efficiently describe the mutations driving cancer in an individual is crucial for precision medicine and can enable targeted therapies explicitly tailored to each patient’s disease (13–16). Determining cancer-driven and cancer-driving mutations in a patient, ideally at single-cell resolution, is challenging. It requires characterizing the millions of cancer-associated mutations, identifying the mutations present in a sample derived from a patient, and distinguishing relevant mutations from technical noise or noncausal biological variation while tracking the cell of origin of mutations.

Several databases have curated large-scale annotated lists of cancer-associated mutations. The Catalogue of Somatic Mutations in Cancer (COSMIC) is one such database, cataloging almost 24 million cancer-associated somatic genomic variants. COSMIC consists of multiple projects, including the cancer mutation census (CMC), whole and targeted genome screens, the cancer gene census, drug resistance screens, and cell lines (17, 18). Each project contains cancer mutations derived from different sources or filtered by varying levels of importance. One group identified 853 pathogenic or likely pathogenic variants from 10,389 cases across 33 cancer types in The Cancer Genome Atlas Program (TCGA) (19). In addition, the Memorial Sloan Kettering Integrated Mutation Profiling of Actionable Cancer Targets (MSK-IMPACT) project has documented 78,066 cancer mutations in 341 key cancer genes (20, 21). The IntOGen meta-database has compiled 257,898,749 mutations across several mutation databases, including TCGA, cBioPortal, and the International Cancer Genome Consortium (ICGC) (22).

Current *de novo* variant detection approaches do not take advantage of these variant databases to improve the sensitivity and interpretability of the variant calling process. Additionally, some existing variant calling tools require significant runtime, on the order of hours to days, for standard sequencing results, and are unable to analyze data from single-cell technologies.

In addition to the creation of variant databases, there has been a steady increase in the amount of genomic sequencing data produced. Approximately 10^21^ bases are sequenced each year, generating 1 million TB of data annually (23). The rate of research studies including next-generation sequencing data is growing rapidly, including studies involving single-cell resolution. Sequencing becomes cheaper each year, with the cost of sequencing a human genome currently costing less than $1,000 (24). Sequencing data is becoming more cost-effective and widespread in a clinical setting, with medicare cost estimates between $500 and $1000 for targeted sequencing assays and $1499.32-3388.18 for whole exome sequencing (25). The increasing adoption of sequencing in the clinic motivates increased accuracy and speed of variant calling in order to allow for real-time diagnosis and treatment planning.

In this study, we introduce varseek, a reference-based variant calling tool that identifies variants with high speed, accuracy, and technology versatility. The varseek program takes as input a reference database of variants and sequencing data from any sequencing technology spanning DNA, bulk RNA, and single-cell RNA sequencing (see Methods, Fig. 1a, Supplementary Fig. 1). We describe the workflow underlying varseek, involving the creation of a reference of variant-containing sequences and the identification of variants to this reference via k-mer matching using kallisto (26–29) followed by filtering. We apply varseek to the analysis of cancer single-cell heterogeneity and population analysis across DNA, bulk RNA, and single-cell RNA (scRNA) sequencing. The varseek tool is open-source, available on GitHub at https://github.com/pachterlab/varseek and PyPi at https://pypi.org/project/varseek.

**Fig. 1.**
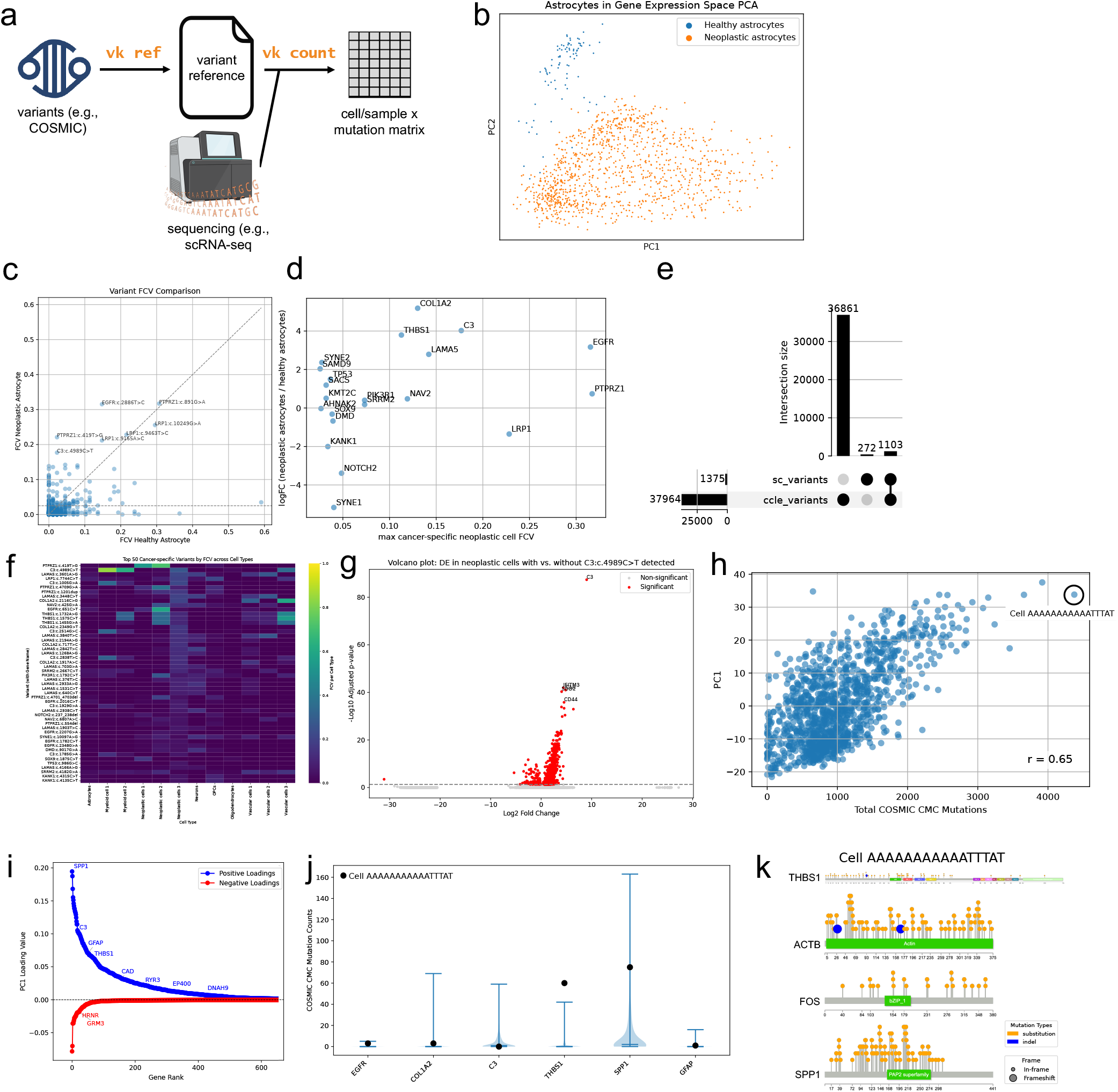
Overview of varseek and its applications to variant calling in single-cell RNA-sequencing data (scRNA-seq). (a) Simple schematic of inputs and outputs of the standard varseek workflow. (b) Principal Component Analysis (PCA) plot in gene expression space of healthy (blue) and neoplastic (orange) astrocytes from a glioblastoma dataset. (c) Variant scatterplot of fraction of cells with variant (FCV) in neoplastic vs. healthy astrocytes. (d) Gene scatterplot of log fold change (logFC) in gene expression between neoplastic and healthy astrocytes vs. the maximum FCV among cancer-specific variants for that gene. (e) UpSet plot of cancer-specific variants as identified by scRNA-seq data compared to bulk RNA-seq data from the cancer cell line encyclopedia (CCLE). (f) Heatmap of FCV for the top 50 cancer-specific variants across the 12 cell types identified by Darmanis et al. (g) Volcano plot of gene expression between neoplastic cells with vs. without the *C3*:4989C>T variant detected. (h) Cell scatterplot of PC1 loading value vs. total COSMIC mutations detected. (i) Loadings of PC1, with some of the genes identified as cancer-specific by cBioPortal labeled. (j) Violin plots of total COSMIC mutations detected across all cells for each gene. Black points = values for the cell circled in black in panel (h). (k) Lollipop plots displaying COSMIC mutations in four selected genes for the cell circled in black in panel (h).

## 2 Results

### varseek identifies mutations at single-cell resolution

To illustrate the ability of varseek to identify mutations at single-cell reoslution, we performed variant calling with varseek on scRNA-seq glioblastoma multiforme (GBM) data generated by Smart-Seq v2 (30) using a variant index genereated from the COSMIC CMC database, which catalogs 5,419,242 mutations associated with cancer (Supplementary Fig. 2). The neoplastic and healthy astrocytes were separated in the 2D principal component analysis (PCA) projection based on gene expression (Fig. 1b). Cancer-specific variants were defined as those present in at least 2.5% of the 1,091 neoplastic astrocytes and in a greater fraction of neoplastic astrocytes than healthy astrocytes. This thresholding identified 5,383 cancer-specific variants, including 73 cancer-specific variants in genes found to be mutated in at least 10 TCGA GBM samples (31) (Fig. 1c). Among these cancer-specific variants were variants in the genes *EGFR* and *SOX9*, which were identified by Darmanis et al. (30) (in their Figure S1D) as strong predictors of the neoplastic phenotype from only their expression. We selected the mutation *C3*:c.4989C>T based on having one of the highest neoplastic fractions of cells with variant (FCV) to perform differential expression analysis (DE) within neoplastic cells. Looking at the genes by differential expression in neoplastic cells and maximum FCV across all variants for each gene revealed *EGFR, C3, COL1A2, THBS1*, and *LAMA5* as genes with cancer-specific variants and cancer-specific expression (Fig.1d).

We also performed variant calling with varseek on bulk RNA-seq GBM data from the Cancer Cell Line Encyclopedia (CCLE) (32). Of the 5,383 cancer-specific variants identified in the GBM single-cell data, 3,719 (69.1%) were also identified in at least half of the 38 CCLE GBM samples (Fig. 1e). Some of the genes with a variant demonstrating high FCV in both scRNA-seq neoplastic cells and CCLE cells include *EGFR*, LRP1, *C3, LAMA5, COL1A2*, and *THBS1* (Supplementary Fig. 3). Among the top 50 cancer-specific variants by FCV in neoplastic cells across all cell types, some variants demonstrate high frequency in myeloid and vascular cells, potentially implicating their role in angiogenesis related to tumor development (Fig. 1f). Comparing the expression of neoplastic cells with the *C3*:c.4989C>T mutation to neoplastic cells without the mutation revealed a strong over-expression of *C3* in mutation-containing cells, followed by the *IFITM3, C1R, SOD2* and *CD44* genes, all known to be associated with GBM proliferation (Fig. 1g).

A varseek analysis enables the characterization of individual cells based on the variants they possess. The total number of COSMIC CMC mutations in a cell had a positive correlation (*r* = 0.65) with the PC1 loading value (Fig. 1h). Some of the genes that contribute the most strongly to PC1 include SPP1, *C3*, GFAP, and *THBS1*, which are involved in glioblastoma processes including hypoxia and tumor microenvironment interaction (Fig. 1i) (33–36). Single-cell analysis revealed an individual cell with both high PC1 and high total number of COSMIC CMC mutations that has a high number of *THBS1* (36) and SPP1 (37) mutations, potentially indicating a more aggressive state (Fig. 1j). Some mutations tend to cluster around hotsopt regions, such as the VWC and TSP_1 domains of *THBS1*, while others are spread more uniformly, such as in FOS (Fig. 1k).

### varseek has high sensitivity at low read coverage

A major advance of varseek is its high sensitivity, including at low coverage. To demonstrate this, we developed a synthetic bulk RNA-seq dataset that represented a range of variant sequencing depths, tumor purities, and variant types (see methods) (Supplementary Fig. 4a). We compared varseek against the most widely used variant calling tools: GATK Haplotype-Caller (4) and Mutect2 (2), Strelka2 (38), VarScan (39), and Deepvariant (6). Although most tools require eight or more reads to call a variant, varseek displays high sensitivity even when there are only 2-3 variant-containing reads at a site (Fig. 2a-c). Furthermore, varseek retains more than 99% sensitivity across all variant sequencing depths for all variant types, while other tools require higher depths for indels than substitutions to reach high sensitivity. Variant allele frequency (VAF) had little to no effect on varseek sensitivity (Fig. 2d-f). All tools had precision >99% at sequencing depth ≥3 reads (data not shown).

**Fig. 2.**
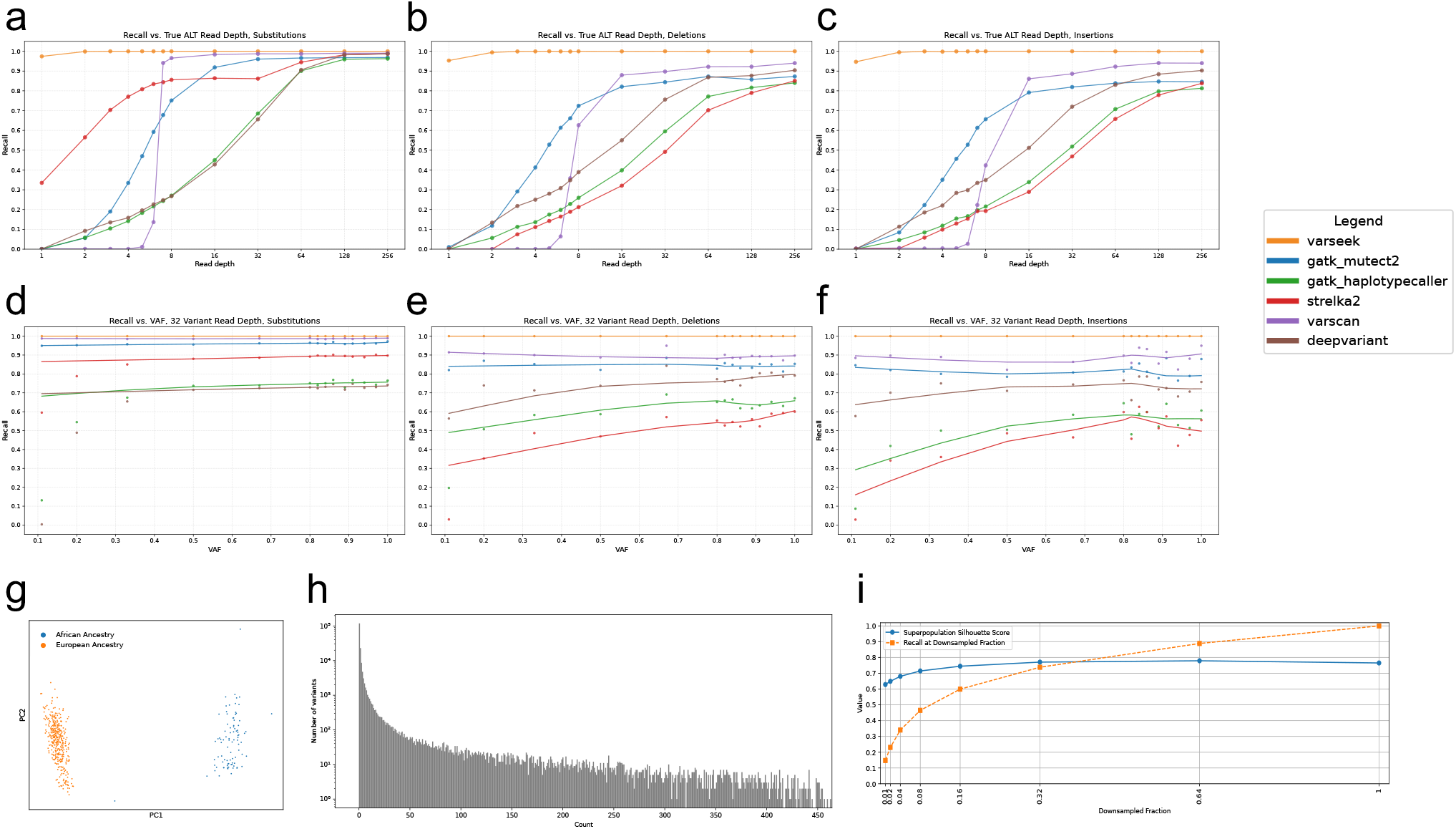
THe varseek method is sensitive in detecting variants. (a-c) Recall vs. variant read depth for substitutions (a), deletions (b), and insertions (c). (d-f) Recall vs. VAF for substitutions (d), deletions (e), and insertions (f) at a fixed variant read depth of 32. (g) Principal Component Analysis (PCA) plot in variant space of African ancestry (blue) and European ancestry (orange) of samples from the Geuvadis dataset. (h) Variant detection histogram corresponding of samples from the Geuvadis dataset. (i) Superpopulation Silhouette Score (blue) and Recall at downsampled fraction (orange) across multiple downsampled fractions of data.

An analysis of all sites with variant sequencing depth ≥ 3 reads revealed that varseek only missed 35 out of 365,497 substitutions, 6 out of 33,196 deletions, and 6 out of 7,415 insertions (Supplementary Fig. 4b-d). Moreover, varseek alone detected 22,517 out of 365,497 substitutions (6.16%), 8,500 out of 33,196 deletions (25.61%), and 4,973 out of 7,415 insertions (67.07%). These results show that while a large percentage of variant-containing reads are correctly aligned (as they are detected by at least one tool based on read alignment), many are not detected at the variant calling step for some tools. To determine the types of indels that most other tools would not detect even at high variant sequencing depth (≥128 reads), we compared variants detected by varseek and at most one other tool to variants detected by varseek and at least two other tools. Some features that were overrepre-sented among poorly detected variants were insertion variants, long variants, and variant location in regions with long homopolymers (Supplementary Fig. 4e-n). Visualization of several deletions, insertions, and duplications confirmed that even when variant calling failed in read alignment-based methods, the associated reads were generally aligned correctly (Supplementary Fig. 5).

Beyond synthetic data, we validated the sensitivity of varseek on the Geuvadis whole exome sequencing (WES) and bulk RNA-seq data in Supplementary Figure 6 (40). We used a previously validated set of 203,850 dbSNP variants that were identified in the transcriptome of at least one Geuvadis sample as our reference variant list, created a reference file using the `vk ref` command, and screened the Geuvadis samples against this reference with `vk count`. As a validation, we selected a random sample and compared the variants detected by varseek in the RNA to the variants detected by previous studies in the DNA. Most variants detected by varseek in the RNA and not detected by other tools in the DNA were either truly present in the RNA according to the read pileup (i.e., either missed by prior DNA variant calling or present as an RNA editing event) or were not truly present in the RNA and were detected in fewer than 5 reads (indicating that a count threshold of 5 would eliminate these false detections). Most variants not detected by varseek in the RNA and detected by other tools in the DNA were either not detected in the RNA according to the read pileup generated by bcftools (i.e., the non-variant allele was expressed) or did not have any reads detected (i.e., the gene was not expressed in RNA at all). We ran `vk count` on the WES and found sizable overlaps between varseek WES and prior DNA variant calling (likely variants not expressed in the transcriptome), as well as between varseek WES and varseek RNA (likely variants missed by prior DNA variant calling) (Supplementary Fig. 6b).

In addition to individual variant detection, we determined the sensitivity of varseek in low-coverage settings based on metrics measuring aggregate information from variants detected. The first such metric is the Silhouette Score, measuring separation of superpopulations (i.e., African ancestry vs. European ancestry) in variant PCA space. On the full dataset, these populations were cleanly separated based on their variants (Fig. 2g). The second such metric is the recall, or the fraction of true variants that were correctly detected (Fig. 2h). We downsampled the sequencing read data at various fractions, finding that the superpopulation Silhouette Score remained relatively consistent between 0.6-0.8 even at 1% of reads sampled, and the recall followed a loosely logarithmic trend (Fig. 2i, Supplementary Fig. 7-8). These results demonstrate the ability of varseek to detect variants and draw meaningful conclusions in low-coverage settings.

### varseek is fast and memory-efficient

We compared the runtime and peak memory use of varseek to the same set of variant calling tools at a range of sequencing data sizes. The runtimes of all the tools increased monotonically with the number of reads. At 256 million reads, the runtimes of all tools in ascending order were as follows: varseek (18 min) < Strelka2 (58 min) < Deepvariant (149 min) < VarScan (681 min) < GATK HaplotypeCaller (804 min) < GATK Mutect2 (881 min) (Fig. 3a). Included in the times of all tools besides varseek is 31.25 minutes for read alignment to the reference genome with STAR. The peak memory usage for varseek, Strelka2, VarScan, and Deepvariant was always below 8GB; the peak memory usage for GATK Haplotype-Caller and GATK Mutect2 was in the range of 17-25GB for datasets of 16+ million reads (Fig. 3b). STAR alignment required a peak memory usage of 35GB. varseek’s low run-time and memory usage can be partially attributed to the optimizations achieved by kallist | bustools (kb), which include the building of a k-mer hash map for quick lookup, the creation of a colored de Bruijn graph as the index for efficient traversal, and the implementation in C (29, 41). The only additional steps performed by varseek during the variant calling step involve FASTQ preprocessing with the fastp library and working with the variant count matrix with the anndata library, both of which have been largely optimized.

**Fig. 3.**
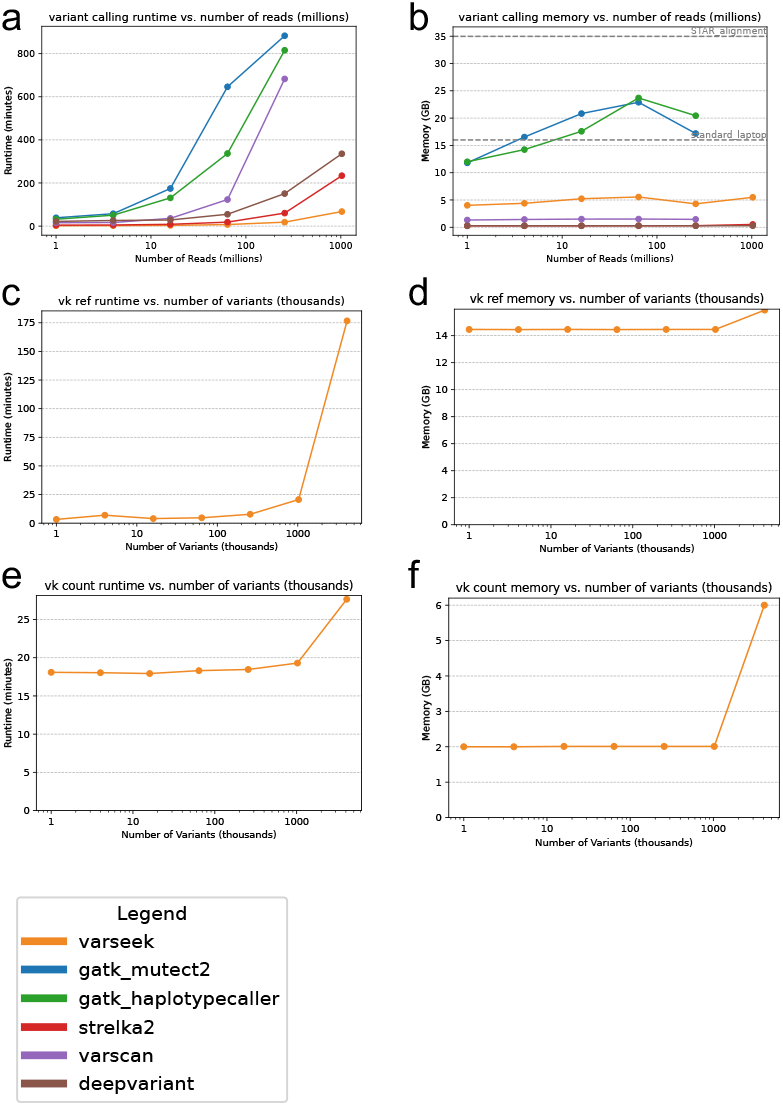
The varseek approach is fast and memory-efficient. (a-b) Variant calling runtime (a) and maximum memory usage (b) of varseek and multiple other variant calling tools by number of reads in the sequencing data. (c-d) `vk ref` runtime (c) and maximum memory usage (d) by number of entries in the variant database. (e-f) `vk count` runtime (e) and maximum memory usage (f) by number of entries in the variant database.

We performed additional analysis on varseek to determine the relationship between variant reference size and run-time and peak memory usage. With a reference containing fewer than 1 million variants, `vk ref` and `vk count` run at a relatively constant runtime with respect to the reference size, with `vk ref` running under 30 minutes and under 16GB RAM, and `vk count` running under 20 minutes and under 2GB RAM (Fig. 3c-f). When the number of variants in the reference surpasses approximately 4 million, run-time increases to approximately 180 minutes for `vk ref` and approximately 30 minutes for `vk count`, and memory increases to approximately 16GB for `vk ref` and 6GB for `vk count`. The increase in runtime of `vk ref` can be mainly attributed to an internal bowtie2 alignment of the variant-containing reference sequence (VCRS) k-mers to the reference genome and transcriptome. As `vk ref` must only be run once per reference database, these runtime costs do not scale with the number of sequencing experiments to be screened.

## 3 Discussion

The varseek method is a reference-based variant caller that enables single-cell resolution variant calling, is adaptable for use with a wide variety of assays and sequencing technologies thanks to its use of kallisto (42), exhibits high sensitivity, including in indel calling, and is efficient in both runtime and memory usage. Using numerous data sets, we have shown that varseek substantially simplifies single-cell genetic analyses, enabling the identification of associations between variants and factors such as cell type, cancer state, and developmental stage.

While k-mers have been utilized to avoid read alignment during variant calling (43), or to better filter (44), varseek benefits from utilizing a novel pseudoalignment-based approach to variant calling that required us to overcome multiple challenges. Most of these relate to false positive matches that can arise when a read shares a k-mer with a VCRS even though the read does not exhibit the corresponding variant. This situation can occur for a multitude of reasons, each of which we address:

1. Sequence similarity between the VCRS and non-variant containing sequence due to flank size (Supplementary Fig. 9a). Solution: We generate VCRSs such that each VCRS contains the variant region and, at most, (k-1) nucleotides flanking each side of the variant, ensuring that each k-mer contains part (if not all) of the variant.
2. Sequence similarity between the VCRS and non-variant containing sequence due to similarity between variant region and variant-flanking regions (indels only) (Supplementary Fig. 9a). Solution: We implemented a flank optimization procedure that shortens the beginning of the left flank based on the degree of overlap between the beginning of the variant sequence and the beginning of the right flank, and analogously that shortens the end of the right flank based on the degree of overlap between the end of the variant sequence and the end of the left flank. For VCRSs where no degree of flank shortening can remove k-mer overlap between the variant and non-variant sequence, we simply remove these VCRSs from the variant reference.
3. Sequence similarity between the VCRS and unrelated regions of the reference genome/transcriptome (Supplementary Fig. 9b). Solution: We filter out VCRSs of low complexity (long homopolymers and/or frequently-repeated nucleotide triplets), as these motifs can be repeated across the genome. Additionally, we align all k-mers of all VCRSs to both the standard genome and transcriptome, and can filter out VCRSs with frequent k-mer matches. As an initial conservative approach with our COSMIC reference, we filter out all VCRSs with any k-mer match to any portion of the human genome or transcriptome. In principle, some VCRSs could be retained and the runtime of `vk ref` could be sped up by providing a BED file for only coding regions of the genome with minimal loss of specificity.
4. Sequence overlap between the VCRS and another close-by variant (Supplementary Fig. 9c). Solution: We used a k > (w+1) to ensure that additional context beyond just the variant region is correctly identified (e.g., k=41, w=37).
5. Sequence overlap between the VCRS and a read due to sequencing errors in low-quality read regions (Supplementary Fig. 9d). Solution: We trim low-quality edges and adaptors off of reads and filter short or low-quality reads to reduce these events.
6. Sequence overlap between the VCRS and another distant variant on another gene (Supplementary Fig. 9e). Solution: We align the reads to the reference transcriptome to determine the gene assignment of each read, and then adjust the variant count matrix by only keeping variant counts where the gene of the variant to which each read maps matches the gene to which each respective read maps.
7. Sequence overlap between the VCRS and a read due to occasional sequencing errors in the middle of high-quality reads (Supplementary Fig. 9f). Solution: Filter out counts from the variant count matrix below a very small minimum threshold (e.g., threshold=3).

False negatives arise when an insufficient number of variant-containing reads possess a k-mer match with the VCRS in the reference index. At the read level, these cases arise when a read contains two additional variants (whether truly occurring in the sample or from sequencing error) flanking the variant of interest on each side within k bases, or when a read contains one additional variant flanking one side of the variant of interest and there are fewer than k bases before the end of the read on the other side of the variant of interest. These false negative cases are minimized by ensuring that k is not set too large with respect to both w and the read length. Approximately 1 in every 250 bases differs between an individual genome and the human reference genome, with a rate of closer to 1 in 1,500 bases differing in coding regions (24). Illumina sequencing technologies have error rates mostly lower than 1 in 1,000 bases (45–48). We assume that *variant*_*frequency*_*rate* = 1*/*1500 uniform (rate of variation from reference genome), *sequencing*_*error*_*rate* = 1*/*1000 uniform (rate of sequencing errors), *k* = 51 (k-mer size), *w* = 50 (VCRS window size), *read*_*length* = 150 (read length), *reads*_*present* = 5 (number of reads with the variant of interest, and *read*_*threshold* = 3 (number of reads needed to detect to call the variant). The probability of missing a read that contains a variant is 0.006-0.033 based on its distance from the read edge. The probability of a false negative (fewer than *read*_*threshold* reads contain a k-mer match) is 0.001-0.033 due to sequencing error alone. Even if there is another variant directly preceding the variant of interest, then the variant of interest can still be detected if the variant of interest is located within the first (*read*_*length*− *k*) bases of the read and if the following k bases do not have sequencing errors for at least *read*_*threshold* reads. The probability of a false negative in this case is 0.271, and goes down to 0.072 if the *reads*_*present* increases from 5 to 7. When building a reference of transcriptome-derived variants, we use the cDNA position (with untranslated regions [UTRs]) rather than the CDS (without UTRs) in order to provide the UTR sequences for added alignment potential. Additionally, for paired-end technologies, to ensure that distinct variants can be detected on each read of a pair, we run the underlying `kb count` command in single parity mode regardless of true data parity. Some of the most important parameters for balancing sensitivity and specificity include w, k, quality_control_fastqs, and qc_against_gene_matrix.

Currently, varseek builds a VCRS index using a single linear reference genome, which limits its ability to detect common variants absent in that reference. By integrating pangenomic approaches (49, 50), one can construct a VCRS for each path through the pangenome graph, thus capturing variation across multiple haplotypes. For variants occurring within *k* bases of each other, a combinatorial indexing strategy could represent all possible combinations of closely spaced variants. When the number of such combinatorial paths becomes large, analyzing co-occurrence patterns across samples may reveal that only a subset of these variant combinations appear together, allowing reduction of the search space. Practically, after running `vk ref` on each path, an auxiliary column could be added to the resulting VCRS metadata dataframe that records the true variant present. This would enable post hoc collapsing of variant count matrix entries following `vk count` corresponding to different pangenome alignment paths that resolve to the same underlying variant, thus streamlining analysis and ensuring consistency in variant interpretation.

Furthermore, our approach, which addresses the challenges above, should be of independent interest as some of the strategies we have employed could be utilized in other read alignment and variant calling algorithms. The varseek algorithms were designed for protocols that involve short reads (10s-100s bp), such that each VCRS, which ranges in length from *k* to (2*k* −1), is comparable in length to the length of the read. This means that there is not substantially more information in a read outside of a region that may overlap with the VCRS. As long-read RNA-seq methods become more common (51), the k-mer-based approaches of varseek could be combined with traditional alignment-based approaches to improve the sensitivity and speed of variant calling while also utilizing the additional context in the reads. Furthermore, varseek may be powerful in combination with pseudoassembly (52), with the latter serving to identify regions from which to construct VCRSs.

While we have showcased varseek’s utility in the applications of cancer characterization, varseek can be applied in a variety of other contexts. Any disease or genetics application for which variants of interest are useful for genotyping, diagnosis, or treatment can benefit from rapid screening with varseek (53–56). For smaller reference databases of variants, varseek will benefit from further speedups as the size of its reference index will be reduced, opening up the possibility for rapid variant calling in clinical settings, a goal that is challenging with current variant calling techniques that rely on all variants to be detected and subsequently screened. Furthermore, varseek can be applied for variant screening in settings of low sequencing depth, including in cell-free DNA (cfDNA) or cell-free RNA screening from liquid biopsies (57–59) and scRNA-seq. This is because varseek identifies variants through k-mer matching, regardless of the depth of the sequencing. varseek can also identify variants equally well in the absence of a paired healthy normal sample. Moreover, varseek can overcome the limitations of variant calling in capturing challenging variants such as gene fusions, splice variants, and RNA-editing events by characterizing the k-mers that distinguish these events from the reference genome.

There is a mature and excellent body of work on sequencing-based cancer detection that we believe can benefit from varseek. For example, Cologuard developed a logistic regression model that combines information regarding *KRAS* mutations, aberrant *NDRG4* and *BMP3* methylation, β*-actin*, and a hemoglobin immunoassay from fecal cfDNA to predict colon cancer (60). More recent methods utilize AI approaches and multimodal data for this task. GRAIL-GALLERI is a multi-cancer screening tool that uses methylation-based analysis of cfDNA in the blood as input to a machine learning model to predict cancer, with a range of effectiveness depending on the cancer type and stage (61, 62). Guardant Health’s Shield test utilizes a combined genomic, epigenomic, and proteomic analysis, demonstrating 83% sensitivity and 90% specificity for detecting advanced colorectal cancer, and 13% sensitivity for advanced precancerous lesions (63). Gencove has shown significant promise in detecting genomic variants at very low coverage (64, 65). Although methods are improving, these applications of sequencing are currently partly limited by the sensitivity or specificity of variant identification.

## 4 Code Availability

- varseek: https://github.com/pachterlab/varseek
- Figures from this study: https://github.com/pachterlab/RLSRWP_2025
- varseek tutorials: https://github.com/pachterlab/varseek-examples

## 5 Data Availability

- COSMIC CMC (17): https://cancer.sanger.ac.uk/cosmic/download/cancer-mutation-census/v101/alldata-cmc
- Glioblastoma Smart-Seq (30): https://www.ncbi.nlm.nih.gov/geo/query/acc.cgi?acc=GSE84465
- CCLE RNA-seq (32): https://www.ebi.ac.uk/ena/browser/view/PRJNA523380
- Geuvadis variants (66): https://www.dropbox.com/s/k9ptc4kep9hmvz5/1kg_phase1_all.tar.gz
- Geuvadis RNA-seq (40): https://www.ebi.ac.uk/ena/browser/view/PRJEB3366
- Geuvadis WES: https://www.internationalgenome.org/data-portal/sample
- Ensembl reference genomes: https://ftp.ensembl.org/pub

## 6 Author Contributions

The varseek project was conceived by JMR, LL and LP. JMR led the development of the varseek algorithms, with the as-sistance of LL and LP. JMR implemented varseek and designed the tests and controls for the method. DKS assisted with kallisto and helped with designing some of the varseek algorithms. RR helped identify applications and interpret results. JMR generated results, made the figures, interpreted the results, and drafted the initial manuscript. All authors read and edited the manuscript.

## 7 Acknowledgments

The varseek project was partially motivated by the kmuxlet idea developed with Sina Booeshaghi, Vasilis Ntranos, and Páll Melsted for demultiplexing single-cell RNA-seq samples using kallisto.

## 8 Methods

### varseek overview

The standard varseek workflow is split into two steps: `vk ref` and `vk count` (Fig. 1a, Supplementary Fig. 1). The purpose of `vk ref` is to create an index of VCRSs to be used as a reference for variant detection; the purpose of `vk count` is to screen sequencing data against this variant index. varseek utilizes the pseudoalignment algorithm from kb, implemented in kb-python, for efficient k-mer matching (29).

The first step, `vk ref`, takes as input a database of variants and the associated reference genome/transcriptome with the variant database, and produces as output an index file of VCRSs. The database of variants can be provided as a list, a variant call format (VCF) file, or a CSV/TSV file with a column for sequence ID and a column for variant information in a format complying with Human Genome Variation Society [HGVS] guidelines (67)). The reference genome/transcriptome can be provided as a list or FASTA file. Each VCRS is designed such that each k-mer (subsequence of length k) of each VCRS contains at least one variant nucleotide while also maximizing sequence length. Trimming and filtering of VCRSs are applied to minimize false positives, and recommended argument values are provided to balance between sensitivity and specificity (more details in the Discussion). The variant index file is generated from a FASTA file containing the VCRSs via kb ref. This index-generation step need only be performed once for a variant database, as the index file can be reused across sequencing experiments.

The second step, `vk count`, takes as input the index of VCRSs produced by `vk ref` and sequencing data, and produces as output a cell/sample x variant matrix. The sequencing data must be provided as a FASTQ file. Trimming and filtering of the sequencing read data are applied, and validation and thresholding of the cell/sample x variant matrix are applied to minimize false positives. As with `vk ref`, recommended argument values are provided to balance between sensitivity and specificity (more details in the Discussion).

### kallisto pseudoalignment

The core of our variant screening methodology consists of the k-mer-based pseudoalignment approach underlying kallisto. To summarize this pseudoalignment approach, kallisto first builds an index from the reference transcriptome consisting of a hash map that maps all k-mers from the reference transcriptome to the position of the node corresponding to that k-mer in a de Bruijn graph for the reference. The kallisto suite of tools also creates a t2g file that maps transcript names to their corresponding gene names. During alignment, kallisto breaks down sequencing reads into k-mers of the same size k, identifies the matching k-mers from the De Bruijn graph, and creates an equivalence class (a set of all viable potential transcript mappings) based on taking the intersection of perfect k-mer matches across all k-mers in the read. The set of transcripts in the equivalence class is then mapped to the unique set of corresponding genes. All ambiguous bases (Ns) in the reads are replaced with As; all ambiguous bases in the reference are replaced with a random base drawn from the uniform distribution. If an equivalence class corresponds to multiple genes, then this read is considered to be multimapped. If the intersection of k-mer matches in a read is the empty set (either if there is not a single k-mer match between a read and the reference de Bruijn graph, or if different sets of k-mers map to different transcripts), then that read is considered to be unmapped. For varseek, by default, we take the variants that correspond to the union, rather than the intersection, of k-mer matches, as this allows variant detection to be independent of which variants comprise the reference file. We balance this by increasing the default k, from 31 for standard kallisto to 51 for standard varseek, as well as providing the option to validate that a read must pseudoalign to the same gene as the gene of the variant to which it maps.

### Variant reference creation and filtering

To build the variant reference, we called `vk ref`, which wraps around `vk build`, `vk info`, `vk filter`, and `kb ref` (Supplementary Fig. 1). Supplementary Figure 2 shows a flowchart that details the variants retained starting from the COSMIC CMC database and ending with the final variant reference used in analysis.

`vk build` takes as input the variant database (as a CSV/TSV file, a VCF file, or a list) and the reference genome from which the variants were annotated. For each variant in the database, it refers to that region of the reference genome, induces the variant, and adds the variant region and *w* nucleotides flanking each side of the variant region to the VCRS FASTA output. The argument value for w must be strictly less than the argument value for k, as this ensures that each k-mer in each VCRS contains the variant. As w decreases in magnitude relative to k, there becomes a greater required overlap on each end of the VCRS to match k-mers during alignment. We set default values for w=47 and k=51. To en-sure that VCRSs do not share k-mers with their WT counterparts, we implement a sequence flank shortening algorithm, wherein the left flank is shortened by the number of nucleotides overlapping between the beginning of the variant region and the beginning of the right flank, and analogously, the right flank is shortened by the number of nucleotides overlapping between the end of the variant region and the end of the left flank. We discarded VCRSs that shared a k-mer with their WT counterpart despite flank shortening, which accounted for <0.001% of sequences in the COSMIC CMC reference and were generally of low sequence complexity. We merged the headers of VCRSs when they were identical to each other or one another’s reverse complement (relevant when the sequencing data is unstranded), as these sequences are indistinguishable in terms of k-mer matching. The entire `vk build` module is entirely vectorized with the pandas Python library.

`vk info` computes statistics regarding the variant reference and stores these in a dataframe, which can be input to `vk filter` to filter the variant reference as desired. The statistics that `vk info` can compute include the following:

- nucleotide composition
- sequence length (mean, total, minimum, maximum)
- whether or not the spliced and unspliced VCRS are identical
- distance to the closest splice junction
- number of unique transcripts and number of unique genes
- the set of nearby variants for each variant
- the number of k-mer overlaps to the normal reference transcriptome and genome
- the number of k-mer overlaps to other VCRSs
- longest homopolymer length
- triplet nucleotide complexity (the number of distinct triplets divided by the number of total triplets)
- whether or not the VCRS is a superstring or a substring of another VCRS

By default, the only computed statistics are (1) the VCRS k-mer alignment and pseudoalignment to the reference genome and transcriptome, (2) longest homopolymer length, and (3) triplet nucleotide complexity.

`vk filter` filters VCRSs from the FASTA file based on a list of provided filters in the format COLUMN:RULE or COLUMN:RULE=VALUE, where COL-UMN is a field in the metadata table, RULE is a rule (valid choices follow), and VALUE is the VALUE to match. Supported rules included numeric comparisons (greater_than, greater_or_equal, less_than, less_or_equal, between_inclusive, between_exclusive), percentile thresholds (top_percent, bottom_percent), equality tests (equal, not_equal, is_in, is_not_in), Boolean checks (is_true, is_false, is_not_true, is_not_false), and nullity (is_null, is_not_null). Only the VCRSs passing all filters will remain in the filtered FASTA file. By default in the `vk ref` pipeline, the applied filters are that only the VCRSs without a k-mer alignment to the reference genome or transcriptome, with five or fewer distinct nucleotide triplets, and with a homopolymer of greater than 10 bases in length.

The filtered VCRS FASTA file is passed into `kb ref` with a custom workflow to generate the VCRS index used for pseudoalignment.

### Variant calling

To screen sequencing data against the variant reference, varseek uses the `varseek count` command, which wraps around `vk fastqpp, kb count, varseek clean`, and `varseek summarize` (Supplementary Fig. 1)

`vk fastqpp` performs preprocessing on the FASTQ files. By default, all processing is turned off. However, if FASTQ process have not been quality controlled, it is recommended to set ‘quality_control_fastqs’, ‘cut_front’, and ‘cut_tail’ all to True. ‘quality_control_fastqs’ enables quality control by applying the fastp package to trim low quality edges and adapters, filter sequences by average base quality and length, and perform additional read processing (68). While fastp is designed for DNA and bulk RNA data, we extend the package to work for barcoded single-cell data by ensuring that read processing only occurs on the file(s) and region(s) of each file with cDNA sequence (as opposed to barcode, UMI, etc.), and that any additional files with barcode/UMI information are adjusted accordingly to ensure synchronization. `vk fastqpp` also includes the argument ‘split_reads_by_Ns_and_low_quality_bases’, which splits reads by Ns and low-quality bases to minimize false alignment due to poor sequencing. This argument is generally not recommended, as it can introduce a high rate of false negatives due to the shearing of reads.

`kb count` pseudoaligns the FASTQ read data to the reference variant index. By default, we use the multimap and union flags, as one read can contain multiple variants, and the detection of any variant in the index should be independent of the detection of any other variant. To ensure that each read in paired-end data can have distinct variants detected, we run `kb count` in single-stranded mode for paired-end data and then correct the resulting matrix file to collapse barcodes across pairs. An additional `kb count` command can pseudoalign the FASTQ read data to the reference genome index if downstream steps require this.

`vk clean` takes as input the Anndata matrix file(s) generated from the kb count step and processes these data. Two of the most important parameters of `vk clean` are ‘qc_against_gene_matrix’ and ‘min_counts’. ‘qc_against_gene_matrix’ cross-references variant alignment by looping through each read, checking its alignment to the variant reference and gene reference, and ensuring that the gene of the variant to which the read maps in the variant reference matches the gene to which the read maps in the gene reference. This step is off by default. ‘min_counts’ performs thresholding on the count matrix to eliminate variant calls at low counts that have a high likelihood of being attributed to sequencing error. `vk clean` also includes a series of parameters to perform standard scRNA-seq processing on the variant matrix based on information from the gene matrix, including cell and gene count thresholding, doublet detection, and normalization, all of which are off by default.

‘vk summarize’ produces a summary text file with some basic analysis on the variant count matrix. Some of these results include the number of variants detected in each cell/sample, the variants detected across the most cells/samples, and the genes with the most variants detected. This step is off by default in the `vk count` command.

### Comparison of varseek to other variant calling tools

All benchmarked variant calling tools required the generation of a BAM file as the first step in the variant calling pipeline. We performed this using STAR with splice-aware settings (69), aligning to GRCh 37, Ensembl release 93 to agree with the COSMIC reference.

The GATK variant calling pipeline followed was the “RNAseq short variant discovery (SNPs + Indels)” best practices guideline (https://gatk.broadinstitute.org/hc/en-us/articles/360035531192-RNAseq-short-variant-discovery-SNPs-Indels). To briefly summarize this process, after performing alignment with STAR, the FASTQ data was converted to an unmapped BAM for input to MergeBamAlignment; MarkDuplicates was run to identify duplicate reads; SplitNCigarReads was was to split reads by alignment gaps over introns; BaseRecalibrator was run to recalibrate the base qualities using information from the alignment and known sites from the 1000 genomes common SNP VCF; and then HaplotypeCaller and Mutect2 were run on the processed BAM file to perform variant calling. All filtering was disabled for each of these tools in order to prevent incorrect filtering that would arise from the synthetic nature of the dataset.

For Strelka2, variant calling was configured in RNA mode (–rna) with default germline workflow parameters. The workflow was set up via configureStrelkaGermlineWorkflow.py and executed locally with multi-threading enabled. For VarScan, we generated pileup files using samtools mpileup with base alignment quality recalculation disabled (-B) and no base quality filtering. Variants were called with VarScan mpileup2cns using minimal filtering thresholds (minimum coverage of 1, at least 1 supporting read, variant frequency ≥1%, and no strand bias filter), outputting VCF format. For DeepVariant, variant calling was performed in accordance with the RNA-seq case study (https://github.com/google/deepvariant/blob/r1.9/docs/deepvariant-rnaseq-case-study.md).

Mosdepth was used to compute per-base coverage of aligned reads, followed by filtering positions exceeding the minimum coverage threshold (set to 3) with bedtools. Variant calling was performed using a downloadable model trained on RNA-seq data, enabling read splitting (split_skip_reads=true) and running with full CPU parallelism to generate VCF outputs.

In order to map the detected variants to the COSMIC database, we converted the COSMIC CMC mutations to a VCF format. We normalized both the COSMIC ground truth VCF and output VCF of each tool with ‘bcftools norm’ (70), and compared VCFs with hap.py (5).

### Synthetic data creation

Synthetic bulk RNA-seq data was created with a series of calls to `vk sim`. We first built a reference of read parents with `vk build` that contained each variant and the 149 nucleotides flanking each side of the variant. We selected reads for each combination of conditions from selections 1-3 below:

- Variant sequencing depth of m=[1, 2, 3, 4, 5, 6, 7, 8, 16, 32, 64, 128, 256]
- WT sequencing depth of w=[0, 1, 2, 3, 4, 5, 6, 7, 8, 16,32, 64, 128, 256]
- [Variant type of substitution, variant type of non-substitution, contains k-mer overlap in the VCRS reference, does not contain k-mer overlap in the VCRS reference, is within 10 bases of a splice junction, is farther than 10 bases from a splice junction]

For each combination of conditions, we randomly selected 350 variant read parents from the COSMIC CMC mutation reference that were not already included in the synthetic data. We then selected m different starting positions between (read_parent_length - read_length + 1), overlapping some starting positions only if m was greater than this value, and created reads of length 150 starting at each point. We repeated a similar procedure for “wildtype” (WT) reads (i.e., non-variant containing reads), drawing from a reference of WT read parents (sequence fragments which had the same start and end points as the mutant fragments but without the variant introduced). We introduced random variants – 85% substitutions, 10% single nucleotide deletions, 5% single nucleotide insertions – in the reads at a rate of 1 in 1000 bases, according to standard Illumina error rates (45–48). We also added 10,000 reads drawn from random regions of the transcriptome not overlapping with any of the COSMIC CMC variants. All base qualities were set to the highest Phred33 score of I.

### Glioblastoma data analysis

We built a reference variant from the COSMIC CMC cDNA with `vk ref` using default arguments. We performed variant calling with `vk count` using standard arguments on Smartseq2 data from Darmanis et al. (30) and bulk RNA-seq data from the CCLE (32) (see Data Availability).

### Geuvadis data analysis

We built a reference variant index from a set of variants previously identified by GATK HaplotypeCaller (see Data Availability). We filtered these variants only to retain those in exon regions, used dbSNP IDs to convert from HGVSg to HGVSc notation, and used the GTF file to convert from CDS to cDNA positions. As the original variant calls were generated by plink, we validated the REF and ALT alleles based on the dbSNP information. We ran `vk ref` on these variants with w=37, k=41, and otherwise default arguments. We performed variant calling with `vk count` using standard arguments (except k=41) on the Geuvadis bulk RNA seq data and whole exome sequencing data (see Data Availability).

### False Negative Calculations

We quantified FN risk under realistic sequencing noise and background variation. For each read, we modeled a flank of length *k*− 1 adjacent to the mutation of interest and treated two independent per-base hazards: (i) a natural background variant with rate *v* and (ii) a sequencing error with rate *s*. For a conservative per-base hazard, we used *p*≈ *v* + *s* (rare-event union bound), so the probability that a flank is clean is (1 −*p*)^*k*−1^. We reported an upper bound on FN by requiring a clean flank on only one side (the case when the variant of interest is on the end of the read), and a lower bound by requiring clean flanks on both sides (the case when the variant of interest is in the middle of the read), i.e., (1 −*p*)^*k*−1^ vs. (1 −*p*)^2(*k*−1)^ (with analogous decompositions using *v* alone or *e* alone to isolate biological variation vs. sequencing error). In the *no linked-variant* scenario, we modeled read count variability by drawing the number of sequenced reads from a binomial distribution; under a worst case, we assumed all informative reads require a single clean flank (one-sided), and under a best case, they require two clean flanks (two-sided). For each draw, we computed the probability of achieving at least a user-specified threshold of “good” reads and summarized FN as the complement. In the *linked-variant* scenario (a second variant immediately upstream), a read is usable only if (a) the focal variant lands in a valid position—probability (read_length − *k*)*/*read_length— and (b) the following *k* − 1 bases are error-free—probability (1 − *s*)^*k*−1^. The per-read variant detection probability is thus

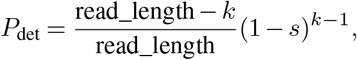

and, given coverage *n*, the probability of observing at least the read threshold is the binomial tail Pr[*X* ≥ *t*], *X* ∼ Binom(*n, P*_det_), with FN equal to 1 − Pr[*X* ≥ *t*].

### Data visualization

Pileup visualizations were performed with the NCBI workbench. Lollipop plots were created with the lollipops package (71). UpSet plots (72) were made with the UpSetPlot package in Python. All other visualizations were made with matplotlib (73) in Python.

