## Supplementary Material for "Reference-based variant detection with varseek"

### Reference-based variant detection with varseek Supplementary Material

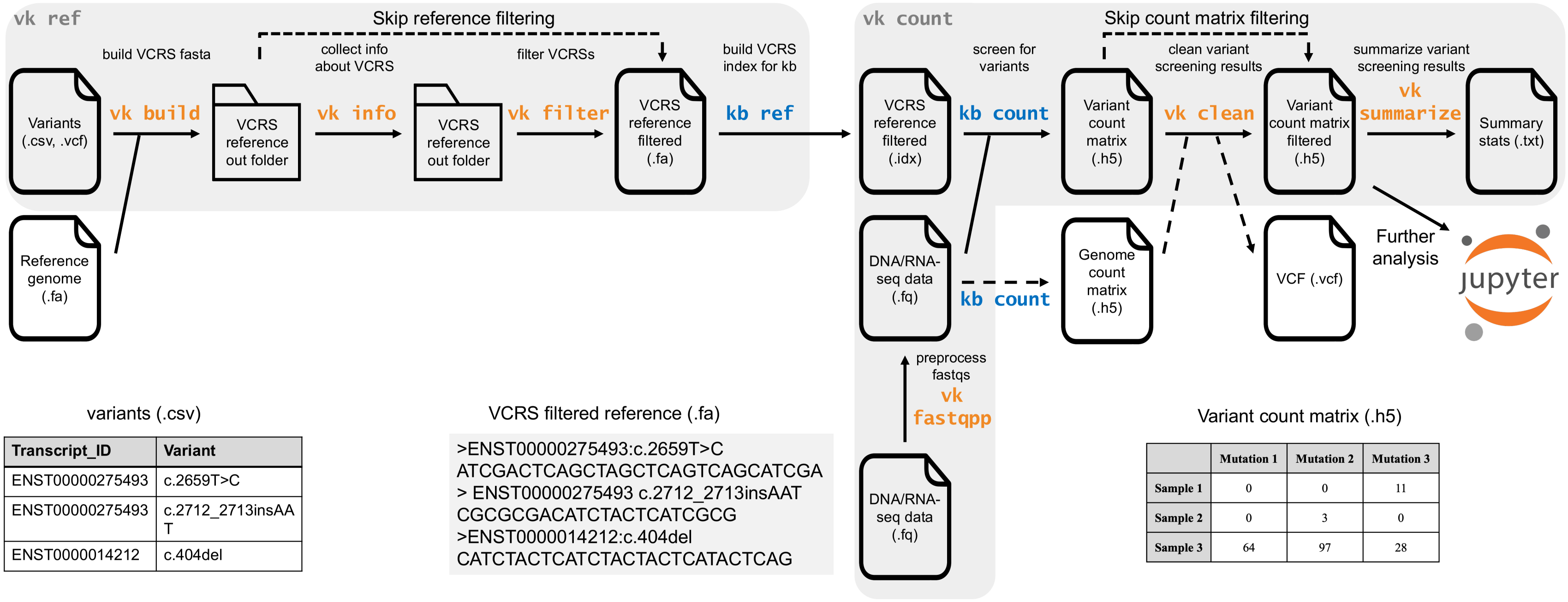


variants (.csv)

| Transcript_ID | Variant |
|---|---|
| ENST00000275493 | c.2659T>C |
| ENST00000275493 | c.2712_2713insAA T |
| ENST0000014212 | c.404del |

VCRS filtered reference (.fa)

```
>ENST00000275493:c.2659T>C
ATCGACTCAGCTAGCTCAGTCAGCATCGA
> ENST00000275493 c.2712_2713insAAT
CGCGCGACATCTACTCATCGCG
>ENST0000014212:c.404del
CATCTACTCATCTACTACTCATACTCAG
```

Variant count matrix (.h5)

| | Mutation 1 | Mutation 2 | Mutation 3 |
|---|---|---|---|
| Sample 1 | 0 | 0 | 11 |
| Sample 2 | 0 | 3 | 0 |
| Sample 3 | 64 | 97 | 28 |

Supplementary Fig. 1: Detailed schematic of varseek. `vk ref` wraps `vk build`, `vk info`, `vk filter`, and kb ref. `vk count` wraps `vk fastqpp`, kb count, `vk clean`, and `vk summarize`. Example file representations of major inputs/outputs are shown below their position in the pipeline. Solid arrows = required inputs/outputs; dashed arrows = optional inputs/outputs.

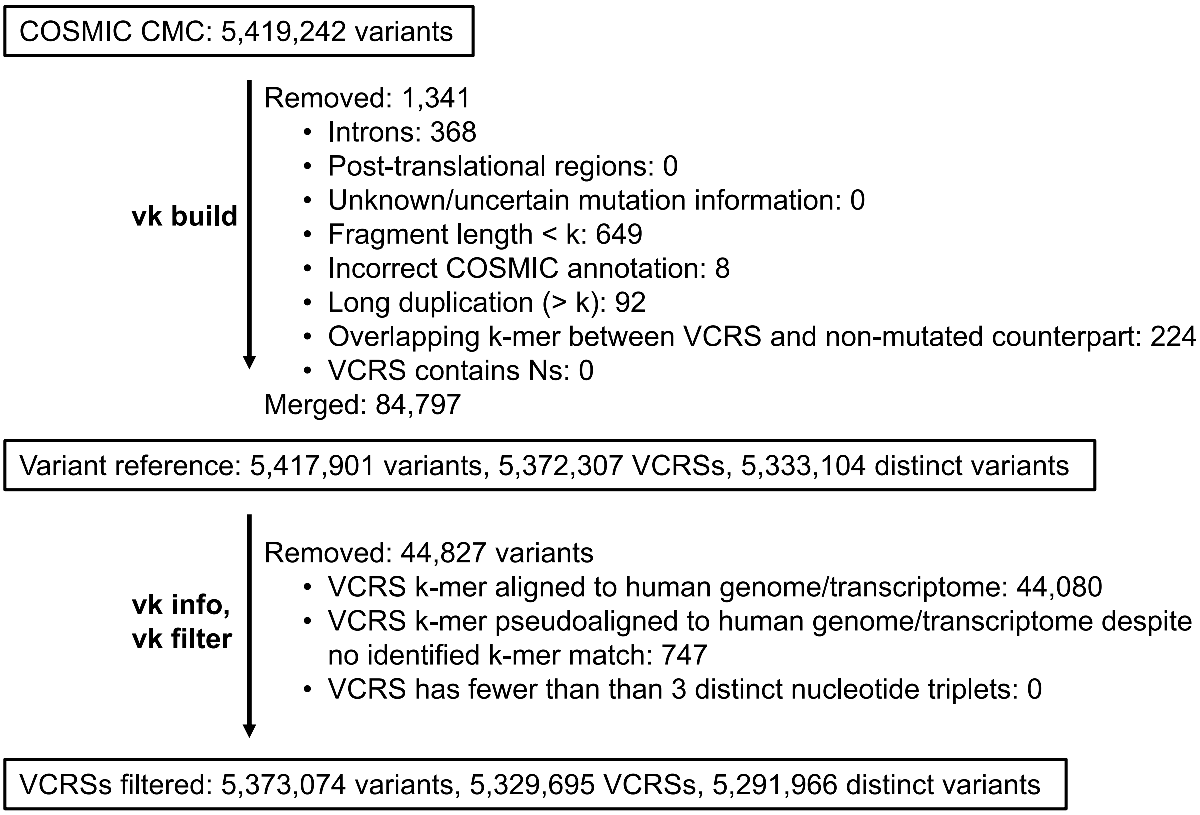


Supplementary Fig. 2: Variant reference index file creation process by `vk ref` from the COSMIC CMC database for analysis of Glioblastoma datasets.

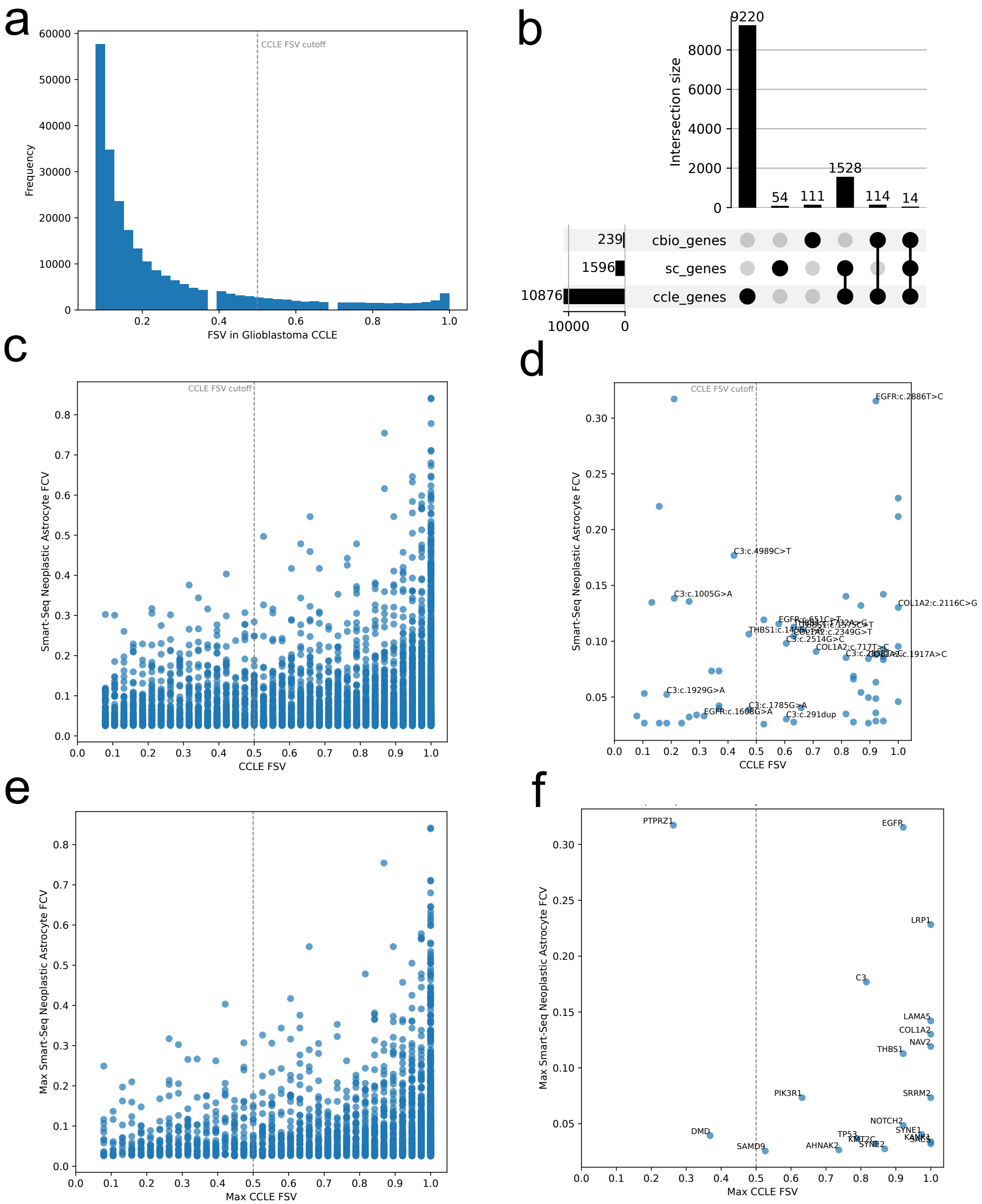

a
CCLE FSV cutoff
Frequency
FSV in Glioblastoma CCLE
b
Intersection size
9220
54
111
1528
114
14
239
1596
10876
cbio_genes
sc_genes
ccle_genes
c
CCLE FSV cutoff
Smart-Seq Neoplastic Astrocyte FCV
CCLE FSV
d
CCLE FSV cutoff
EGFR:c.2886T>C
C3:c.4989C>T
C3:c.1005G>A
COL1A2:c.2116C>G
THBS1:c.1453T>C
COL1A2:c.2349G>T
C3:c.2514G>C
COL1A2:c.717T>C
C3:c.1929G>A
C3:c.1785G>A
EGFR:c.1608G>A
C3:c.291dup
Smart-Seq Neoplastic Astrocyte FCV
CCLE FSV
e
Max Smart-Seq Neoplastic Astrocyte FCV
Max CCLE FSV
f
PTPRZ1
EGFR
LRP1
C3
LAMA5
COL1A2
NAV2
THBS1
PIK3R1
SRRM2
NOTCH2
SYNE1
DMD
TP53
KMT2C
KANK1
SAMD9
AHNAK2
SYNE2
Max Smart-Seq Neoplastic Astrocyte FCV
Max CCLE FSV

Supplementary Fig. 3: Additional glioblastoma analysis between single-cell RNA-seq (scRNA-seq) data and Cancer Cell Line Encyclopedia (CCLE) bulk RNA-seq data. (a) Histogram of fraction of samples with variant (FSV) for all variants in CCLE data. FSV=0.5 used as a threshold for denoting a cancer-specific variant in CCLE data (dashed gray line). (b) UpSet plot comparing the sets of cancer-specific genes as identified by cBioPortal TCGA (mutated in at least 10 samples), scRNA-seq data (containing a mutation that is present in at least 2.5% of neoplastic cells and in a higher percentage of neoplastic cells than healthy cells), and CCLE (FSV $\geq 0.5$). (c-d) Scatterplot for variants of scRNA-seq neoplastic fraction of cells with variant (FCV) vs. CCLE FSV. (c) is for all variants, and (d) is for cancer-specific variants (cBio definition). (e-f) Scatterplot for genes of scRNA-seq neoplastic cells max FCV across all variants for the gene vs. CCLE max FSV across all variants for the gene. (e) is for all genes, and (f) is for genes of cancer-specific variants (cBio definition).

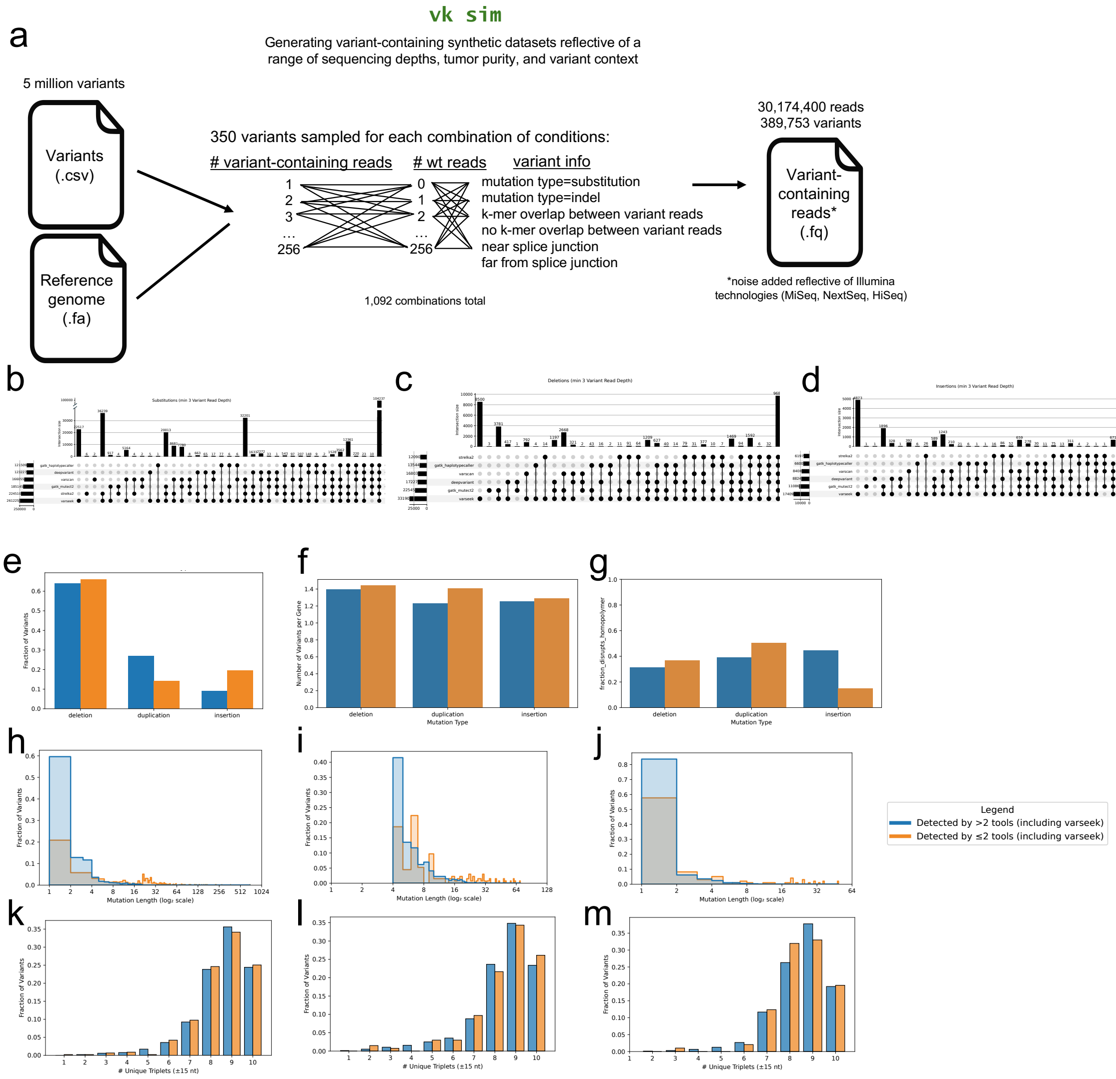

vk sim
Generating variant-containing synthetic datasets reflective of a range of sequencing depths, tumor purity, and variant context
5 million variants
Variants (.csv)
Reference genome (.fa)
350 variants sampled for each combination of conditions:
### variant-containing reads
### wt reads
variant info
mutation type=substitution
mutation type=indel
k-mer overlap between variant reads
no k-mer overlap between variant reads
near splice junction
far from splice junction
1,092 combinations total
30,174,400 reads
389,753 variants
Variant-containing reads* (.fq)
*noise added reflective of Illumina technologies (MiSeq, NextSeq, HiSeq)
Legend
Detected by >2 tools (including varseek)
Detected by ≤2 tools (including varseek)
Mutation Length (log₂ scale)
### Unique Triplets (±15 nt)
Fraction of Variants
Number of Variants per Gene
Mutation Type

Supplementary Fig. 4: Creation of synthetic RNA-seq dataset and analysis of variant detection. (a) Schematic of the inputs and outputs to the function `vk sim` to produce the dataset. Each combination of conditions in # variant-containing reads, # wt reads, and variant info had 350 variants drawn from it. (b-d) UpSet plot of detection of substitution (b), deletion (c), and insertion (d) Variants with a minimum variant read depth of 3 across all tools. (e-m) Analysis of indels with a minimum variant read depth of 128, stratified by those detected by varseek and at most one other tool (orange) or varseek and at least two other tools (blue). (e) Fraction of variants in each variant type. (f) Average number of variants per gene. (g) Fraction of variants that disrupt a homopolymer (at least three identical bases in a row). (h-j) Variant length for deletions (h), insertions (i), and duplications (j). (l-n) Local sequence complexity (defined by the number of unique triplets in a reading frame within 15 nucleotides of the variant) for deletions (l), insertions (m), and duplications (n).

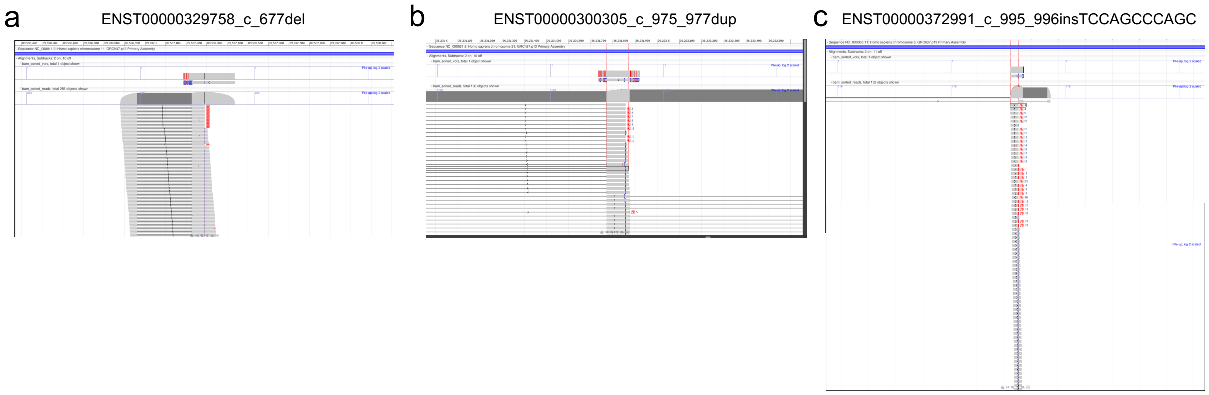


Supplementary Fig. 5: Visualization of the pileups of three variants detected by only varseek and at most one other tool. (a) ENST00000329758:c.677del (a deletion). (b) ENST00000300305:c.975_977dup (a duplication). ENST00000372991:c.995_996insTCCAGCCCAGC (an insertion). Note that positions are given relative to the cDNA of each transcript.

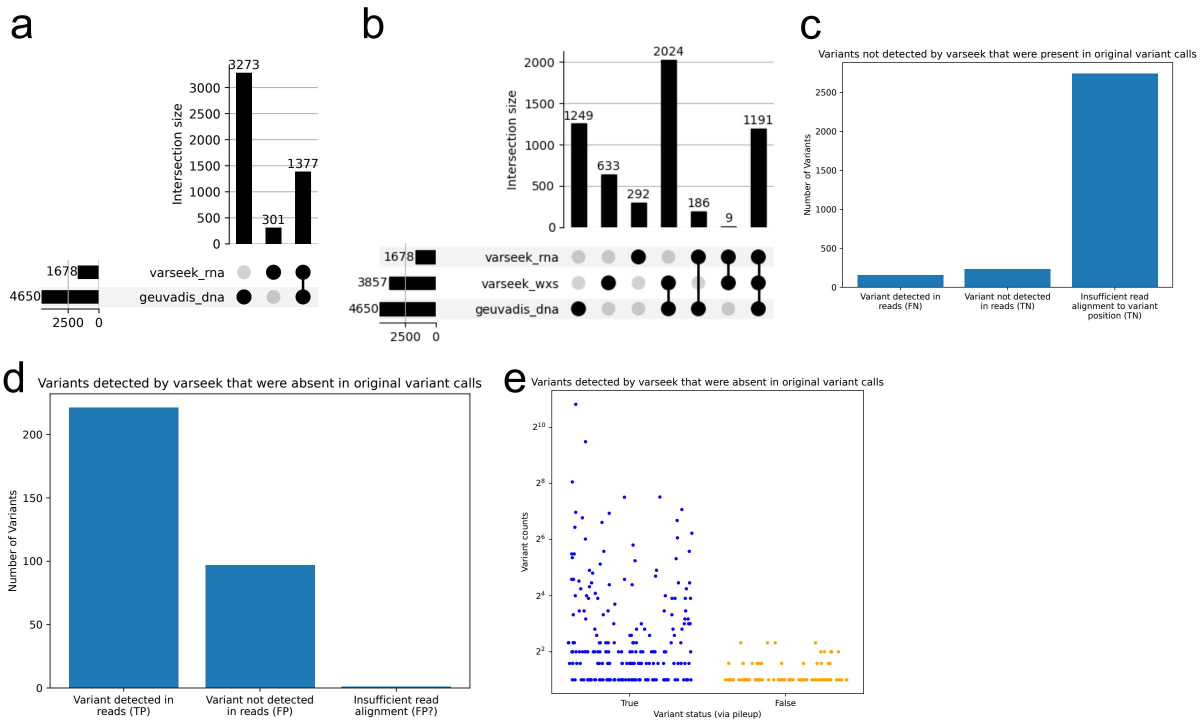


Supplementary Fig. 6: Validation of varseek on a sample from the Geuvadis dataset. (a) UpSet plot comparing variants detected by varseek in bulk RNA-seq to variants detected from GATK HaplotypeCaller in gene-coding regions of whole genome sequencing. (b) UpSet plot similar to (a) but with the addition of an entry for varseek applied to whole exome sequencing. (c) Explanation for cases where varseek did not detect a read and GATK HaplotypeCaller did detect a read. (d) Explanation for cases where varseek detected a read and GATK HaplotypeCaller did not detect a read. (e) Stratification by number of reads of variants undetected by varseek but detected by both GATK HaplotypeCaller and pileup.

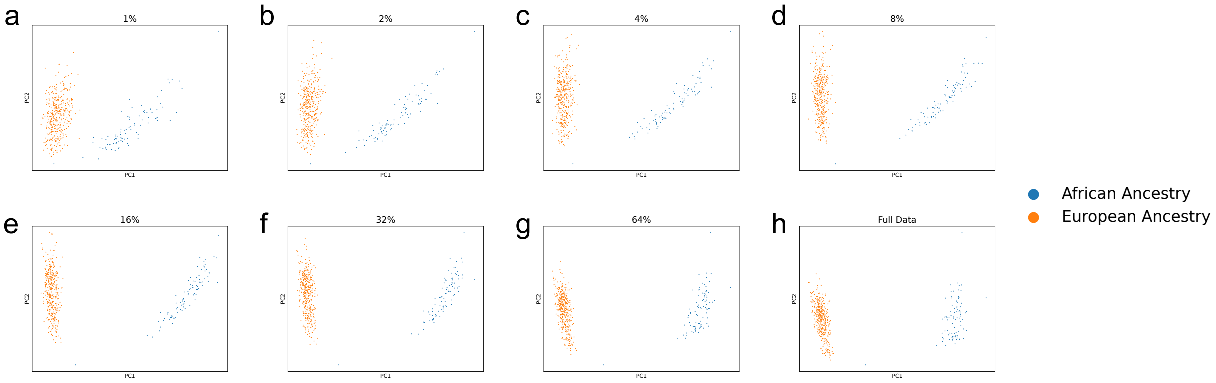


Supplementary Fig. 7: PCA plots of Geuvadis data colored by superpopulation at different downsampled fractions corresponding to Fig. 2i. Each percent provides the amount of data remaining (i.e., 1 - percentage downsampled). (a) 1%. (b) 2%. (c) 4%. (d) 8%. (e) 16%. (f) 32%. (g) 64%. (f) 100% (full data).

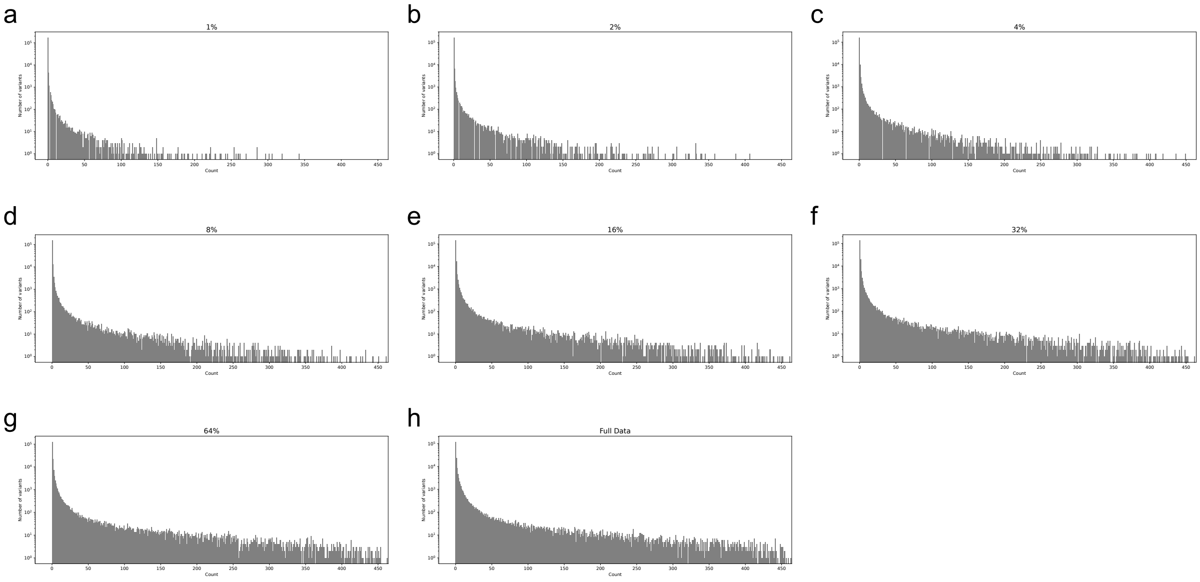


Supplementary Fig. 8: Variant detection histogram corresponding to Fig. 2i. Each percent provides the amount of data remaining (i.e., 1 - percentage downsampled). (a) 1%. (b) 2%. (c) 4%. (d) 8%. (e) 16%. (f) 32%. (g) 64%. (f) 100% (full data).

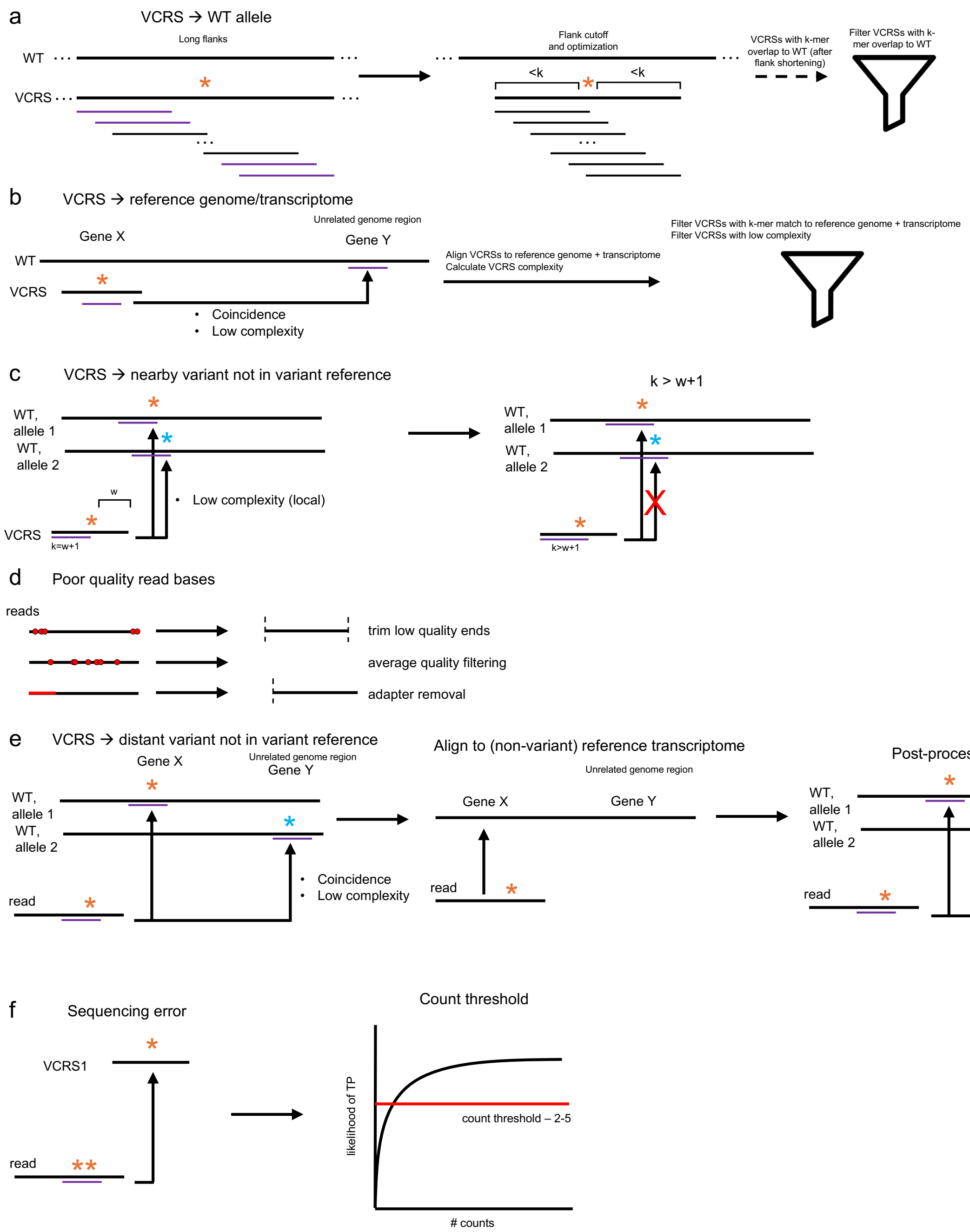

a
VCRS → WT allele
Long flanks
WT
VCRS
Flank cutoff and optimization
<k
<k
VCRSs with k-mer overlap to WT (after flank shortening)
Filter VCRSs with k-mer overlap to WT
b
VCRS → reference genome/transcriptome
Gene X
Unrelated genome region
Gene Y
WT
VCRS
Coincidence
Low complexity
Align VCRSs to reference genome + transcriptome
Calculate VCRS complexity
Filter VCRSs with k-mer match to reference genome + transcriptome
Filter VCRSs with low complexity
c
VCRS → nearby variant not in variant reference
WT, allele 1
WT, allele 2
w
Low complexity (local)
VCRS
k=w+1
k > w+1
k>w+1
d
Poor quality read bases
reads
trim low quality ends
average quality filtering
adapter removal
e
VCRS → distant variant not in variant reference
Gene X
Unrelated genome region
Gene Y
WT, allele 1
WT, allele 2
Coincidence
Low complexity
read
Align to (non-variant) reference transcriptome
Post-processing with BUS file
f
Sequencing error
VCRS1
read
Count threshold
likelihood of TP
count threshold – 2-5
### counts

Supplementary Fig. 9: Methods implemented in varseek for controlling for false positives. All solutions are described in more detail in the Discussion and Methods sections. (a) Case when a k-mer from the variant containing reference sequence (VCRS) is shared with a k-mer in the non-variant containing sequence. (b) Case when a k-mer from the VCRS is shared with a k-mer from another part of the reference genome or transcriptome. (c) Case when a k-mer from the VCRS is shared with a nearby variant. (d) Case when a low-quality read region erroneously contains a VCRS k-mer due to sequencing error. (e) Case when a k-mer from the VCRS is shared with a distant variant. (f) Case when a high-quality read region erroneously contains a VCRS k-mer due to sequencing error.
